# Multiplex mapping of protein-protein interaction interfaces

**DOI:** 10.1101/2025.01.28.635392

**Authors:** Jingxuan He, Ling-Nan Zou, Vidhi Pareek, Stephen J. Benkovic

**Author notes:** These authors contributed equally to this work (Jingxuan He, Ling-Nan Zou). **Corresponding:** Jingxuan He, Ling-Nan Zou, **Email:** (Jingxuan He), (Ling-Nan Zou). **Author Contributions:** JH, LNZ, and SJB conceived the study. JH and LNZ designed, conducted, and analyzed SpARC-mapping experiments. JH prepared materials for in vivo photocrosslinking, VP ran the protein mass spectrometry, VP and LNZ analyzed the mass spectrometry data. JH and LNZ wrote the manuscript. **Competing Interest Statement:** Penn State has filed a provisional patent on SpARC-map and related technologies with JH, LNZ, and SJB as inventors.

## Abstract

We describe peptide mapping through *Sp*lit *A*ntibiotic *R*esistance *C*omplementation (SpARC-map), a method to identify the probable interface between two interacting proteins. Our method is based on in vivo affinity selection inside a bacterial host, and uses high throughput DNA sequencing to infer the probable protein-protein interaction (PPI) interfaces. SpARC-map uses only routine microbiology techniques, with no reliance on specialized instrumentation, dedicated reagents, or reconstituting protein complexes in vitro. SpARC-map can be tuned to detect PPIs over a broad range of affinities, multiplexed to probe multiple PPIs in parallel, and its nonspecific background can be precisely measured, enabling the sensitive detection of weak PPIs. Using SpARC-map, we recover known PPI interfaces in the p21-PCNA, p53-MDM2, and MYC-MAX complexes. We also use SpARC-map to probe the purinosome, the weakly bound complex of six purine biosynthetic enzymes, where no PPI interfaces are known. There, we identify interfaces that satisfy structural requirements for substrate channeling, as well as protein surfaces that participate in multiple distinct interactions, which we validate using site-specific photocrosslinking in live human cells. Finally, we show that SpARC-map results can impose stringent constraints on machine learning based structure prediction.

**Significance Statement:** Protein-protein interactions (PPIs) are vital to biological function, and the identity of the PPI interface can lend valuable insights on the structure/function of protein complexes. Identifying PPI interfaces using standard structural biology approaches, such as x-ray protein crystallography or cryo-electron microscopy, is technically challenging and require reconstituting protein complexes in vitro. As an alternative, SpARC-map is a sensitive and potentially high throughput method for identifying probable PPI interfaces, requiring only standard laboratory methods and straightforward bioinformatic analyses. Its accessibility gives SpARC-map the potential to be a tool of first resort for identifying PPIs and probable PPI interfaces, thereby facilitating scientific discovery across molecular and cellular biology.

## Introduction

In living cells, protein-protein interactions (PPIs) are vital to biological function. Identifying PPIs, and elucidating the composition/structure of protein complexes, are core tasks in molecular and cellular biology research. Here, the location of the PPI interface can illuminate the structure of the protein complex, reveal the functional consequences of the interaction, and point toward cellular mechanisms that regulate it (1, 2). PPI interfaces are also attractive drug targets, and disrupting disease-relevant PPIs is a promising approach towards precision medicine (3–6). Yet, most PPIs interfaces remain unknown. In large part, this is because structural biology tools such as x-ray protein crystallography and cryo-electron microscopy have significant instrumental/technical demands that limit their accessibility; furthermore, these tools require protein complexes to be reconstituted in vitro, which can be challenging, and their throughput is low. Advances in machine learning (ML) based structural prediction (7–10) raises the possibility that where experimental structures are absent, computation models can fill in the gap. However, compared to the large dataset of protein structures available to train ML models, the number of protein complexes with solved structures is far smaller; consequently, the accuracy of ML-based predictions for protein complex structures requires further evaluations grounded in experimental data.

Here, we describe peptide mapping via split-antibiotic resistance complementation (SpARC-map), an indirect method for mapping the probable interface between two interacting proteins. It uses in vivo affinity scanning inside a bacterial host, read out via high throughput DNA sequencing, to infer the location of PPI interfaces. SpARC-map employs only routine microbiological methods, without reliance on specialized instrumentation or dedicated reagents. It does not require reconstituting protein complexes in vitro, is tunable to detect PPIs over a broad range of binding affinities, and can be multiplexed to probe multiple PPIs in parallel. The nonspecific background of SpARC-map can be precisely measured and characterized, allowing its users to make a rigorous and quantitative distinction between specific and nonspecific interactions, thereby enabling the detection of weak, but still specific, interactions.

As a proof of concept, we show SpARC-map can recover known PPI interfaces of human p21-PCNA, p53-MDM2, and MYC-MAX complexes. As a further challenge, we use SpARC-map to map, in multiplex, probable PPI interfaces in the human purinosome, the multi-enzyme complex responsible for de novo purine biosynthesis (11). This weakly-bound complex has never been reconstituted in vitro, in part or in toto, and no PPI interfaces are known (12). Using SpARC-map, we identify probable PPI interfaces that satisfy structural demands for substrate channeling (13); multispecific protein surfaces that participate in multiple PPIs, which we validate using site-specific photocrosslinking in live human cells; and interfaces decorated with post translational modifications (PTMs) that may regulate purinosome assembly. Our results demonstrate SpARC-map can identify true PPI interfaces in even weakly bound protein complexes, to reveal biologically significant insights into their structure, function, and regulation.

## Results

### Peptide mapping through split-antibiotic resistance complementation

SpARC-map is motivated by the observation that many PPIs are mediated by small linear or structural motifs, or are anchored by a small number of hot spots (14–18). The concept is simple: we scan, piecewise, the amino acid sequence of a protein (prey), to identify prey peptide fragments that show affinity for its interaction partner (bait); we expect these peptides will correspond to protein domains that contain the bait-prey interface (Fig. 1A). While a prey peptide fragment is unlikely to adopt same structure as the corresponding region on the intact prey protein, so long as its conformational ensemble overlaps that of the (local) protein structure, the bait-prey peptide interaction can mimic the bait-intact prey interaction (19–21). However, the binding affinity will be reduced due to the entropic penalty of locking in prey peptide conformation upon its binding to the bait.

**Figure 1.**
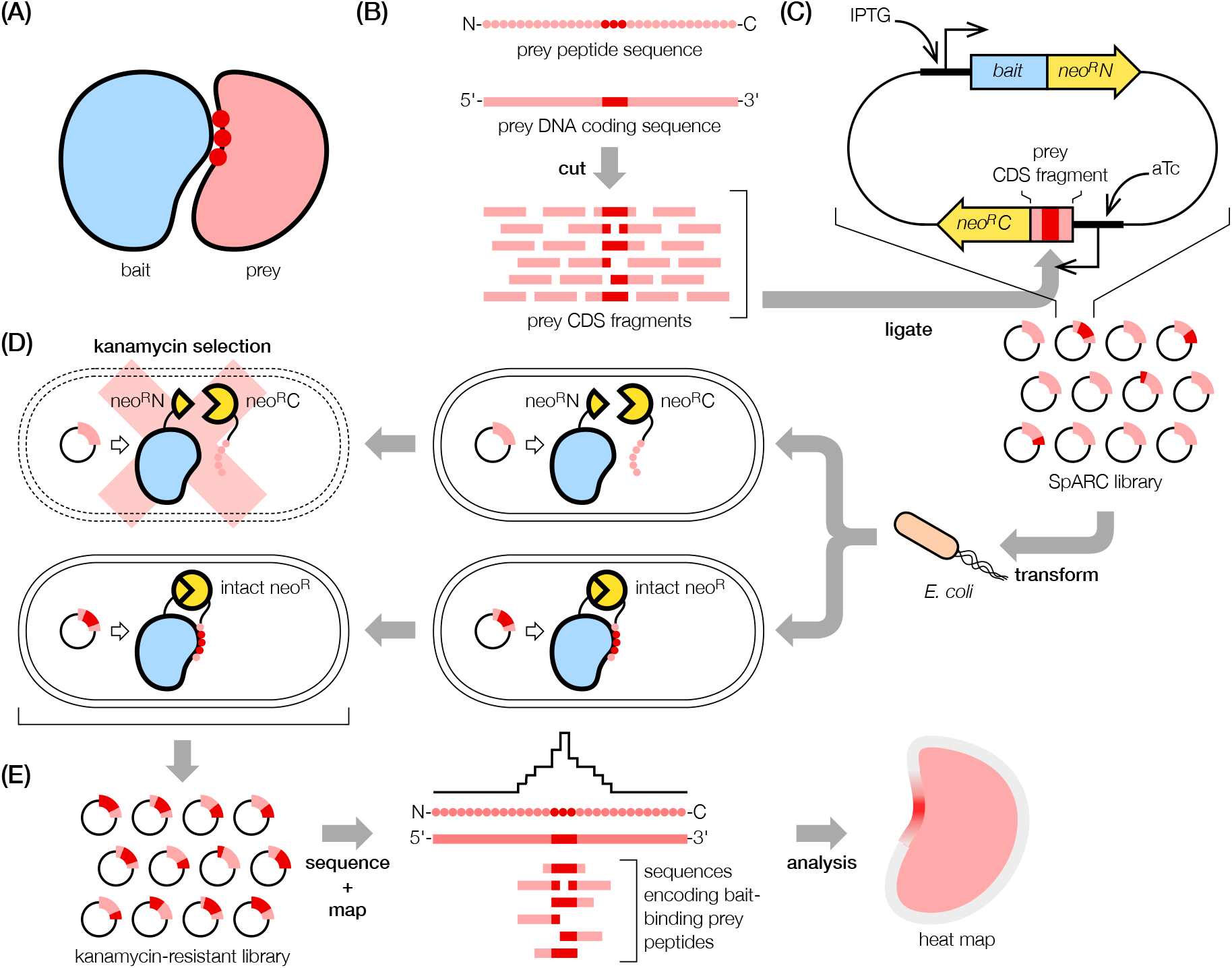
Schematic of SpARC-map. **(A)** Pair of interacting bait and prey proteins; prey residues on the bait-prey PPI interface are highlighted. **(B)** dsDNA containing the prey CDS is sheared into random fragments and **(C)** ligated into the bicistronic SpARC vector, in upstream fusion with the *neo*^*R*^*C*. The intact bait protein is fused to *neo*^*R*^*N*; the fusion terminus depends on the bait. Expression of the bait and prey peptide are under the control of ITPG and aTc inducible promoters. **(D)** The SpARC library is transformed into *E. coli* host and selected in the presence of kanamycin. SpARC vectors that express bait-binding prey peptides can result in kanamycin resistance through the reconstitution of neo^R^N and neo^R^C fragments into a functional neo^R^. **(E)** SpARC vectors from kanamycin-resistant colonies are sequenced, mapped, and analyzed to extract the PPI interface.

We begin by constructing a library of bacterial expression vectors (SpARC vectors); these contain a bicistronic cassette expressing both the bait protein and a small prey peptide (Fig. 1C). The prey peptide coding sequences (100-300 bp) are generated by shearing dsDNA containing the prey protein coding sequence (CDS) (Fig. 1B). To affinity-select bait-binding prey peptides, we engineered host antibiotic resistance conditional on bait-prey peptide binding. We split aminoglycoside phosphotransferase (neo^R^, UniProt P00552), which inactivates kanamycin, into neo^R^N (AAs 1-59) and neo^R^C (AAs 59-264) fragments. Neither alone can inactivate kanamycin, but when fused to interacting proteins, neo^R^N and neo^R^C fragments can reconstitute a functional neo^R^ (22). By fusing the bait and the prey peptide respectively to neo^R^N and neo^R^C, SpARC vectors expressing bait-binding prey peptides can endow their host with kanamycin resistance (Fig. 1D) — if the binding affinity is sufficiently high.

SpARC-map selection is tunable. In the SpARC vector, bait and prey peptide expressions are controlled by orthogonal inducible promoters: IPTG (isopropyl-β-D-thiogalactopyranoside) inducible for bait, aTc (anhydrotetracycline) inducible for prey peptide (Fig. 1C). High bait and/or prey peptide expression favors complexation, enabling even weak interactions to reconstitute sufficient neo^R^ activity to result in kanamycin resistance; conversely, low bait and/or prey peptide expression means only strong bait-prey peptide interactions can lead to kanamycin resistance. By adjusting concentrations of kanamycin and inducers, the selection threshold of SpARC-map can be tuned over a range of bait-prey peptide affinities; we estimate a maximum dynamic range for the apparent bait-prey peptide dissociation constant of *K*_D_^app^ = 10 nM - 1 mM (Supplementary Materials). Post selection, we identify probable bait-binding regions on the prey using next-generation sequencing (NGS) to profile SpARC vectors from resistant bacteria (Fig. 1E).

### Identifying the non-specific background

To distinguish between specific and nonspecific bait-prey peptide interactions, we note that if the background signal of nonspecific interactions, defined as the binding between the bait and *random* peptides (23), can be precisely measured and modelled, then given the measurement of a bona fide peptide fragment of the prey, we can rigorously evaluate whether its interaction with the bait is nonspecific. Prey peptides that are unlikely to be non-specific binders are presumably specific interactors that contain (portions of) the bait-prey PPI interface. Fortunately, constructing the SpARC library automatically supplies a pool of random DNA sequences, encoding random peptides, with which we can determine the bait-random peptide interaction background. Because prey peptide coding sequences are generated by shearing dsDNA containing the full prey CDS, only 1/18 of resulting dsDNA fragments will ligate in-sense and translate into a prey peptide fused in frame to neo^R^C. The remainder will be inverted, and/or frameshifted, and/or contain extra bases modulo 3. Most such nonsense inserts will disrupt in-frame fusion with neo^R^C on 3’ of the peptide coding sequence, or contain premature stop codons; SpARC vectors with such inserts cannot result in kanamycin resistance (Fig. 2A). However, some nonsense inserts will translate through to effectively random peptides fused to neo^R^C; these can interact with the bait and potentially reconstitute neo^R^. We can recognize such random peptides in the NGS results, and for each species *i*, quantify its abundance *n*_*i*_. The distribution *P*_*NS*_(*n*_*i*_) is the expected background due to nonspecific bait-random peptide binding.

**Figure 2.**
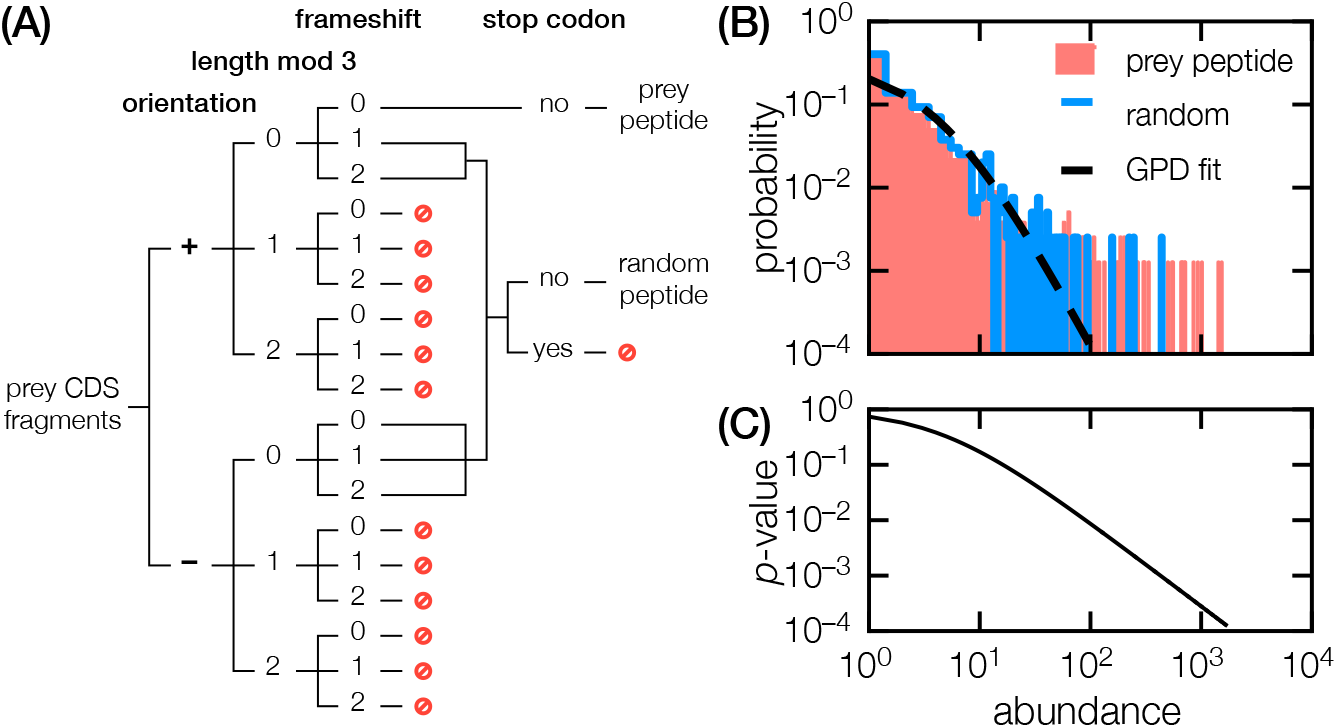
Analysis of NGS species abundance to detect non-specific interactions. **(A)** In addition to in-sense prey CDS fragments (orientation = +, length mod 3 = 0, frameshift = 0), some nonsense fragments, when ligated into the SpARC vector upstream of *neo*^*R*^*C*, can translate into a random peptide fused in frame with neo^R^C fragment. **(B)** From a representative SpARC-map dataset: measured abundance of NGS species encoding random peptides (blue line) and the resulting GPD fit (black dashed line), compared against NGS species that encode true prey peptides (red shaded). **(C)** From the GPD fitted to the random peptide abundance, we can compute a *p*-value for any bona fide prey peptide given its abundance; those with small *p* are unlikely to be nonspecific binders.

For a random peptide to endow its host with sufficient kanamycin resistance to survive selection, its bait-binding affinity must exceed the threshold determined by the selection condition: greater the exceedance, greater the survival. The Pickands–Balkema–De Haan Theorem, a central result of Extreme Value Theory, states the distribution of threshold exceedances, under generous conditions, follows a Generalized Pareto Distribution (GPD) (24). We therefore model the measured non-specific background as *P*_NS_(*n*_*i*_) = GPD(*n*_*i*_; *µ, σ, ξ*), with parameters *µ, σ*, and *ξ* (Fig. 2B) (25). Then, given a NGS species encoding a bona fide prey peptide, we can use the modelled nonspecific background *P*_NS_(*n*_*i*_) to evaluate, by computing appropriate *p*-values, how well its abundance can be explained by bait-random peptide binding (Fig. 2C). Prey peptides whose abundances are consistent with random binding should be excluded from delineating the probable PPI interface.

### Recovering known PPI interfaces in structurally resolved complexes

As a proof-of-concept, we sought to use SpARC-map to identify PPI interfaces in three well-characterized heteromeric human protein complexes: p21/CDKN1A (Uniprot P38936) and PCNA (UniProt P12004); p53 (Uniprot P04637) and MDM2 (Uniprot Q00987); MAX (Uniprot P61244) and MYC (Uniprot P01106). All three complexes have been structurally resolved using x-ray protein crystallography, and their respective PPI interfaces are known.

#### p21-PCNA

PCNA forms a homo-trimeric ring-shaped DNA clamp that is essential for DNA replication and repair; p21, an inhibitor of cyclin-dependent kinases, can bind PCNA via its C-terminal PIP-box motif (p21 AAs 140-164) and inhibit normal PCNA functions (26–30). Figure 3A shows SpARC-map affinity heatmaps (using p21 and PCNA as baits) for the p21-PCNA interaction, displaying the probability a given prey residue is tiled by a mapped NGS read. On p21, we observe a hotspot spanning p21 AAs 139-160, corresponding to the true p21 interface with PCNA. From the tiling by distinct p21 fragments, we can infer while p21^140-164^ has the highest PCNA affinity (is most abundant), truncation to p21^140-158^ still retains PCNA-binding but at a lower affinity (has lower abundance, Supplementary Data 1). This is consistent with known PCNA binding affinities of p21 peptides: *K*_*D*_ = 6.4±2.8 nM for p21^140-163^, and 67±9 nM for p21^143-157^ (31). For PCNA, we observe a hot spot on AAs 104-139, in good agreement with the crystal structure of PCNA in complex with p21^139-160^, where the main PCNA interface with p21 is on PCNA AAs 118-131 (32).

**Figure 3.**
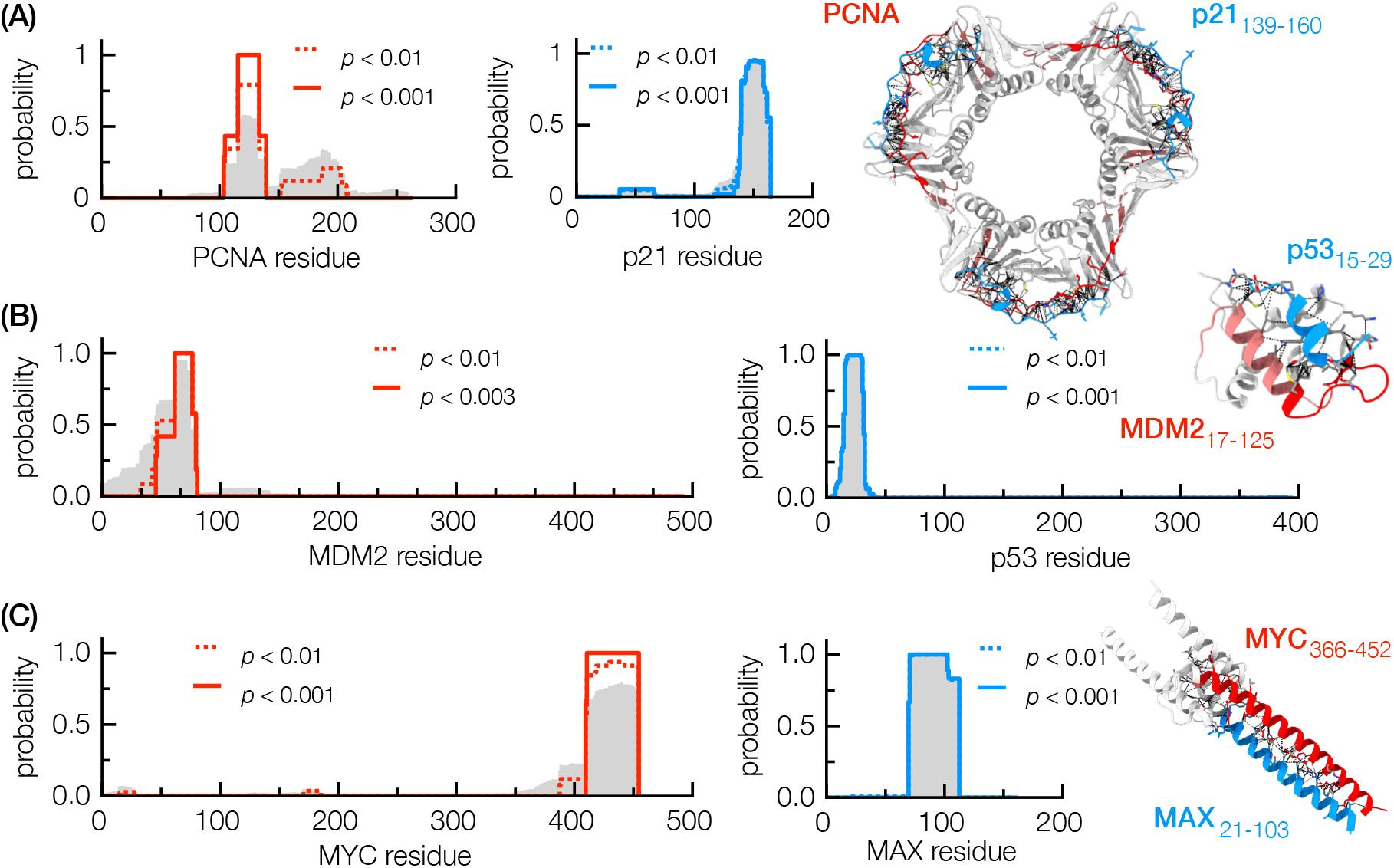
SpARC-map recovers known PPI interfaces of structurally revolved protein complexes. Affinity heatmaps compiled from total NGS reads unfiltered by statistical significance are shown in gray fill; heatmaps compiled from statistically significant (as indicated by *p*-values) NGS species are shown in dotted and solid lines. **(A)** Heatmaps for the p21-PCNA interaction, shown alongside with the crystal structure of PCNA homo-trimer in complex with three p21^139-160^ peptides (PDB 1AXC). PCNA is colored by its affinity heatmap for p21 (red/white = most/least probable); p21^139-160^ peptides are shown in blue. **(B)** Heatmaps for the p53-MDM2 interaction, shown alongside the crystal structure of MDM2^17-125^ in complex with the p53^15-29^ peptide (PDB 1YCR). MDM2^17-125^ is colored by its affinity heatmap for p53 (red/white = most/least probable); p53^15-29^ peptide is shown in blue. **(C)** Heatmap for MAX-binding on MYC interaction, shown alongside the crystal structure of MAX^21-103^ in complex with MYC^366-452^ (PDB 6G6K). MYC^366-452^ is colored by its affinity heatmap for p53 (red/white = most/least probable); MAX^21-103^ is colored by its affinity heatmap for MYC (blue/white = most/least probable).

#### p53-MDM2

p53 is a transcription factor and a critical tumor suppressor; its mutations are among the most common mutations found in human cancers (33, 34). p53 interacts with the E3 ubiquitin ligase MDM2, a key cellular antagonist of p53 activity, via its transactivating domain on p53 AAs 12-27; the reciprocal p53 interacting region on MDM2 is on MDM2’s N-terminal SWIB/MDM2 domain (AAs 25-109) (35). Reported binding affinities for this interaction ranges from *K*_*D*_ = 70-700 nM (36). SpARC-map (using p53 and MDM2^18-125^ as baits) heatmaps for this interaction display statistically significant hot spots on p53 AAs 19-29 and MDM2 AAs 46-80 (Figure 3B), in good agreement with the PPI interface from the crystal structure of p53^15-29^ in complex with MDM2^17-125^ (37).

#### MAX-MYC

MYC is a basic helix-loop-helix/leucine zipper (bHLH/LZ) family transcription factor that has important roles in cellular differentiation and proliferation (38); elevated and persistent MYC expression is a hallmark of many cancers (39). MYC heterodimerizes with MAX, another bHLH/LZ transcription factor, to form sequence-specific DNA-binding complex (40, 41). This interaction is mediated via MYC’s and MAX’s respective bHLH/LZ domains (MYC AAs 369-449, MAX AAs 23-102), and has an affinity in the range *K*_*D*_ = 200-600 nM (42). In SpARC-map (using MAX and MYC as baits) heatmaps, we find a hot spot spanning MYC AAs 411-454, encompassing portions of MYC’s bHLH (AAs 367-421) domain and its entire LZ (AAs 428-448) domain; while on MAX, we observe a hot spot on MAX AAs 70-102, encompassing the C-terminus of MAX’s bHLH domain and the entirety of its LZ domain. These results are in good agreement with the PPI interface found in the crystal structure of MYC^366-452^ in complex with MAX^21-103^ (43).

### Mapping PPI interfaces in the purinosome

Compared to p21-PCNA, p53-MDM2, and MYC-MAX, the human purinosome is far more complicated protein complex. It consists of six enzymes — PPAT (amidophosphoribosyltransferase, UniProt Q06203), GART (trifunctional glycineamide ribonucleotide synthetase/formyltransferase/phosphoribosylaminoimidazole synthetase, UniProt P22102), PFAS (phosphoribosylformylglycinamidine synthase, UniProt O05167), PAICS (bifunctional phosphoribosylaminoimidazole carboxylase/phosphoribosylaminoimidazole succinocarboxamide synthetase, UniProt P22234), ADSL (adenylosuccinate lyase, UniProt P30566), and ATIC (bi-func– tional 5-aminoimidazole-4-carboxamide ribonucleotide formyltransferase/inosine monophosphate cyclohydrolase, UniProt P31939) — responsible for de novo purine biosynthesis (DNPB, Figure S1) (11, 12). Imaging studies have shown these enzymes can colocalize into punctate bodies in purine-starved cells, and multiple molecular studies have found evidence for PPIs (44–49). However, it is unknown if any of these PPIs are direct. All evidence indicates the purinosome is a weakly-bound — it has never been reconstituted in vitro, wholly or partially, and no PPI interfaces are known.

To map probable PPI interfaces in the purinosome, we constructed and screened six multiplexed libraries, one per each purinosome enzyme as bait; all of them have as prey CDS fragments from all six enzymes in multiplex (Supplementary Materials, Table S1). We focus here on probable PPI interfaces between sequential enzymes. Given the clear metabolomic signatures of substrate channeling in the purinosome, we expect at least some enzyme pairs that catalyze sequential reactions should physically interact in such a manner as to bring up- and downstream enzyme domains and/or active sites into proximity, to facilitate direct substrate transfer between them.

#### PPAT-GART

PRA (5-phosphoribosylamine), product of PPAT in reaction 1 of DNPB, is the substrate of GART’s GAR (glycineamide ribonucleotide) synthase domain for reaction 2. The instability of PRA (half-life <1 minute under physiological conditions) was early motivation for suggesting PPAT and GART must physically interact to transfer such a labile intermediate (50, 51). On GART, we found probable PPAT binding sites on AAs 14-55 (*p*<0.001) and 159-232 (*p*<0.003); these flank the GAR synthase active site that receives PRA from PPAT (Fig. 4A), as expected from the structural requirements substrate channeling. On PPAT, the probable GART-binding site is on its C-terminus, AAs 468-504 (*p*<0.001) (Fig. 4B).

**Figure 4.**
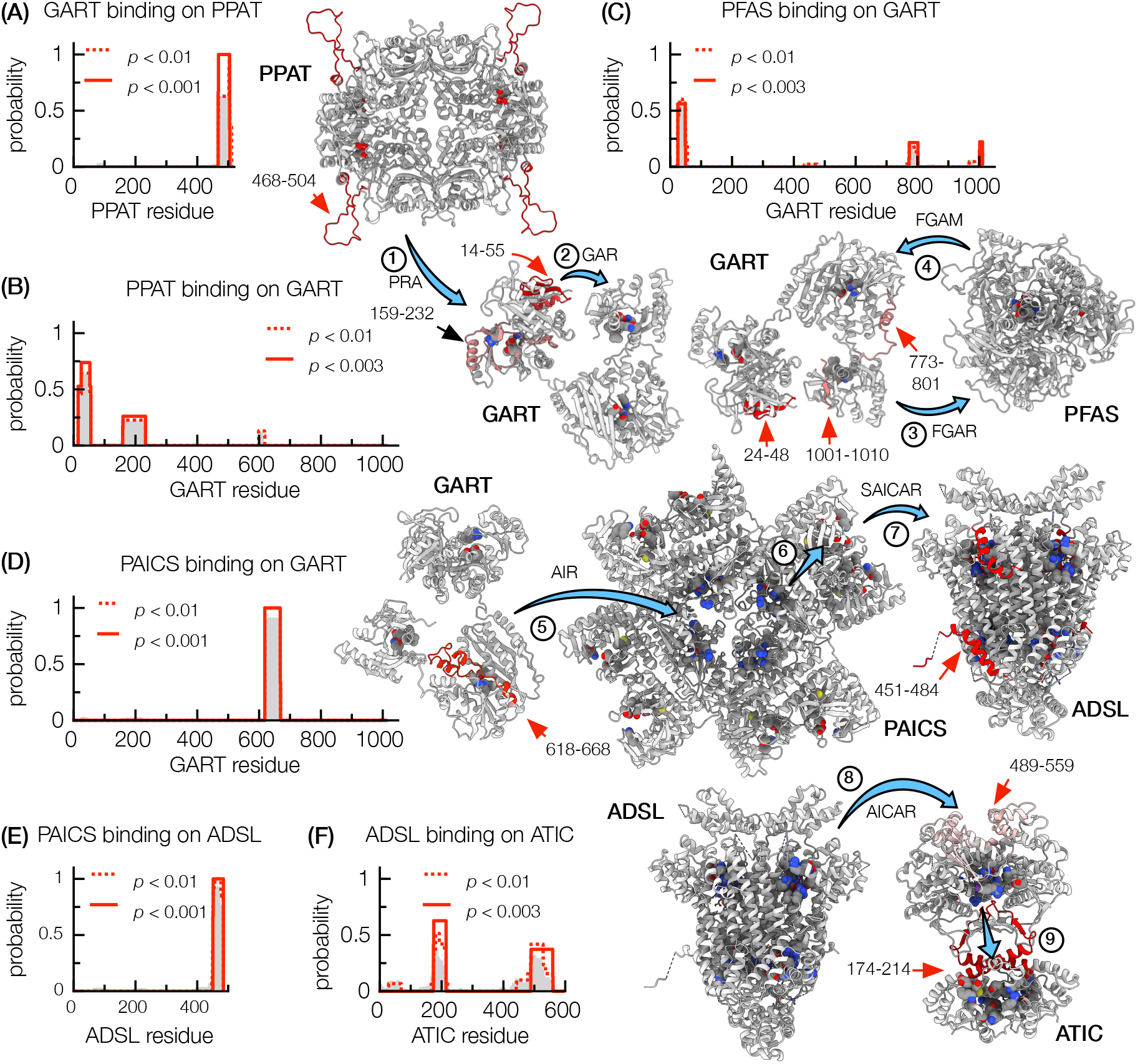
High confidence (*p*<0.003) PPI interfaces between sequential enzymes in the purinosome. Protein structures are colored by affinity heat map (red/white = most/least probable). Spherical atoms highlight enzyme active sites and ligand binding sites. Blue arrows indicate the direction of substrate transfer from upstream to downstream enzyme (domain). Heatmap plots show probable **(A)** GART (monomer) binding sites on PPAT (homo-tetramer); **(B)** PPAT binding sites on GART; **(C)** PFAS (monomer) binding sites on GART; **(D)** PAICS (homo-octamer) binding sites on GART; **(E)** PAICS binding sites on ADSL (homo-tetramer); **(F)** ADSL binding sites on ATIC (homo-dimer). Heatmaps compiled from total NGS reads unfiltered by statistical significance are shown in gray fill.

#### GART-PFAS

GART catalyze reactions 2, 3, 5 of DNPB, while PFAS catalyzes reaction 4, receiving its substrate FGAR (phosphoribosyl-N-formylglycineamide) from GART’s GAR transformylase domain, and sending its product 5-phosphoribosylformylglycineamidine (FGAM) forward to GART’s AIR (5-aminoimidazole ribotide) synthase domain. On GART, we found three probable PFAS-binding sites (*p*<0.003): AAs 24-48, AAs 773-801 on the AIR synthase domain that receives FGAM from PFAS, and AAs 1001-1010 on the GAR transformylase domain that supplies FGAR to PFAS (Fig. 4C); the latter two are structurally consistent with substrate channeling from GART to PFAS and back. We could not identify the reciprocal GART-binding interface on PFAS.

#### GART-PAICS

AIR, the product of GART in reaction 5, is the first substrate of PAICS, which catalyzes reactions 6 and 7. On GART, we found a probable PAICS binding site, AAs 618-669 (*p*<10^−3^), on the AIR synthase domain that supplies AIR to PAICS (Fig. 4D), structurally consistent with substrate transfer from GART to PAICS. We could not identify the reciprocal GART-binding interface on PAICS.

#### PAICS-ADSL

SAICAR (phosphoribosylaminoimidazolesuccinocarboxamide), the product of PAICS in reaction 7, is the substrate for ADSL for reaction 8. On ADSL, we found a probable PAICS binding site on AAs 451-484 (*p*<0.001) (Fig. 4E). On PAICS, we identified three potential ADSL-binding interfaces, AAs 251-270, 281-348, and 413-425, with low confidence (*p*<0.004 – 0.01); two of these, AAs 251-270 and 413-425, are located on PAICS’s SAICAR synthase domain that supplies ADSL.

#### ADSL-ATIC

ADSL supplies AICAR (5-aminoimidazole-4-carboxamide ribonucleotide) as the initial substrate for ATIC, which catalyzes reactions 9 and 10. On ATIC, we found probable ADSL binding site on AAs 174-214 (*p*<0.002) and 489-529 (*p*<0.003); the latter lines the active site entrance on ATIC’s AICAR transformylase domain that receives AICAR from ADSL (Fig. 4F), and is structurally consistent with substrate channeling from ADSL to GART. We identified two low confidence (*p*<0.008) ATIC-binding sites on ADSL, AAs 73-130 and 447-484; the latter coincides with the probable PAICS-binding site on ADSL.

In addition to expected PPI interfaces between sequential enzymes, we also found probable PPI interfaces between enzymes that are distal along the DNPB pathway (Supplemental Materials, Table S2). Indeed, we found a dense network of probable PPI interfaces (at least partially), indicative of a direct physical interaction, connecting every enzyme pair. Surprisingly, some enzymes appear to use the same surface regions to participate in multiple PPIs. For example, ADSL appears to use the same C-terminal region (AAs 451-484) to interact with distal enzymes PPAT and PFAS, as well as enzymes that catalyze immediate upstream (PAICS) and downstream (ATIC) reactions; however, the probable GART-binding site on ADSL (AAs 391-402) is distinct.

### Validating PPI interfaces via in vivo photocrosslinking

While some of the probable purinosome PPI interfaces identified by SpARC-map are expected (being consistent with structural requirements of substrate channeling), others (e.g. the multispecific C-terminal region of ADSL) were surprising. We therefore sought to validate SpARC-map identifications using site-specific photocrosslinking in live human cells. To do so, we use genetically encoded unnatural amino acid incorporation to insert *p*-azido-L-phenylalanine (azF) at defined sites on a protein (52–54). Upon UV-activation, the aryl azide group on azF can attack nearby double bonds, primary amines, C-H and N-H groups, etc., to form a covalent bond. Thus azF, when inserted into the surface of a bait protein, can photocrosslink prey proteins that interact with the bait via that surface (Fig. 5A); its small reactive radius (<0.3 nm) and short lifetime (~1 ns post-photoactivation) minimizes nonspecific crosslinking due to random molecular collisions.

**Figure 5.**
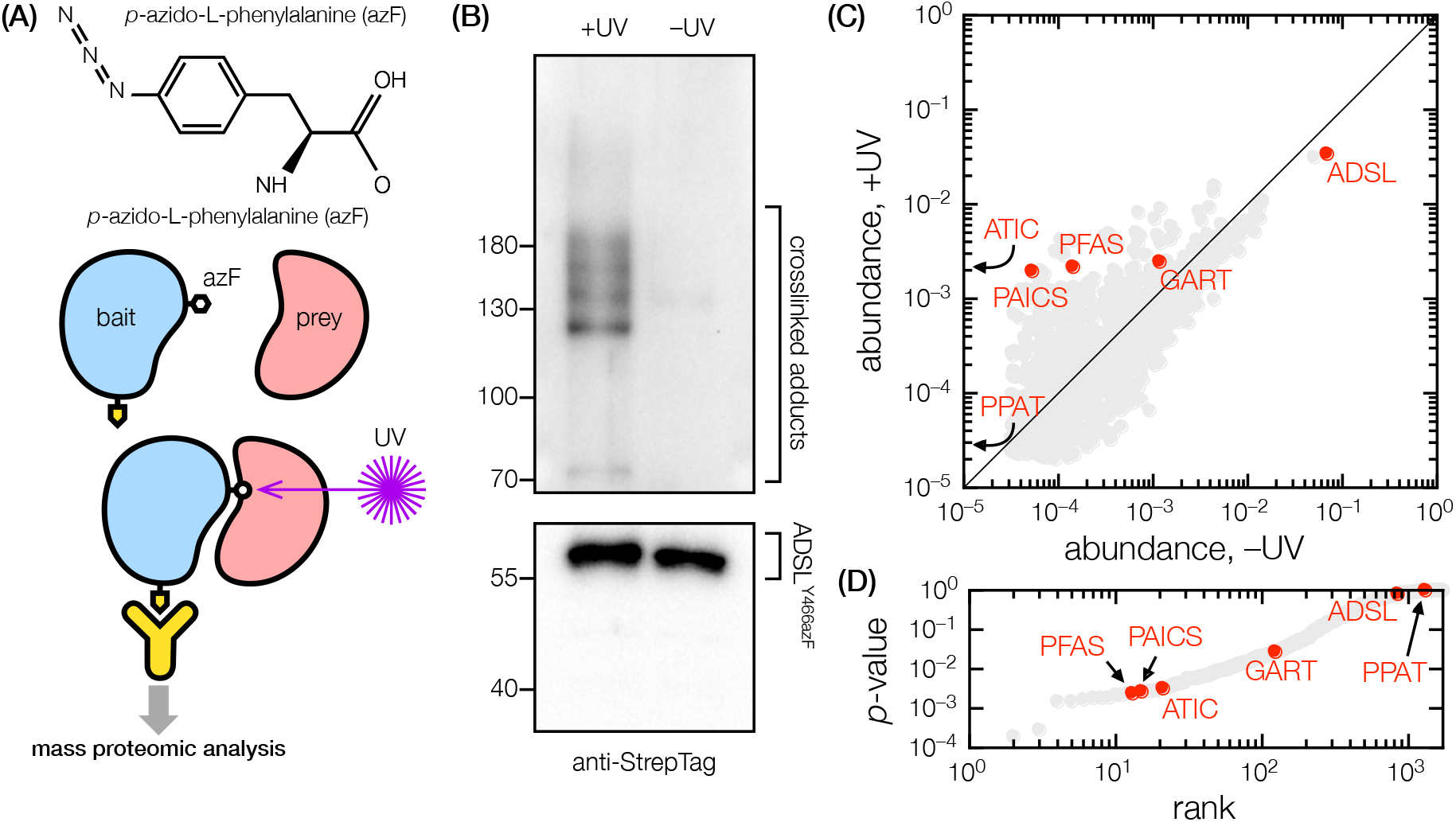
Site-specific photocrosslinking with ADSL^Y466azF^ in live cells. **(A**) Photoactive *p*-azido-L-phenylalanine (azF) is inserted onto a PPI interface of a bait protein. Upon UV-irradiation, azF can form a covalent crosslink between the bait and any prey that interact with the bait via the specific interface. The resulting adduct is pulled down and analyzed. **(B)** Western blot showing anti-StrepTag pull-down from HEK293T transfected with 2×StrepTag-ADSL^Y466azF^. High molecular weight crosslinked adducts are apparent following UV-irradiation. **(C)** Mass proteomic analysis of protein abundance in anti-StrepTag pull-downs with and without UV irradiation. The six purinosome enzymes are highlighted in red. **(D)** *p*-values in ascending rank, indicating PFAS, PAICS, and ATIC are significantly enriched in UV-irradiated vs non-irradiated samples.

We focused on validating ADSL^451-484^ as the multispecific PPI interface between ADSL and other purinosome enzymes. We constructed a mutant ADSL containing the substitution Y466azF, as well as a 2×StepTag epitope tag fused to its N-terminus, and expressed it in HEK293T cells cultured in purine-depleted media. Following brief UV irradiation (for photocrosslinking samples), we lyse the cells and pull-down ADSL^Y466azF^ (plus its crosslinked adducts and co-precipitants) for analysis using mass spectrometry (Fig. 5B). Pulled-down proteins should include those that interact specifically with ADSL but not via the C-terminal region, and those that interact with ADSL specifically via ADSL^451-484^. Abundance of the latter, but not the former, should be enhanced by UV-irradiation.

Of the four DNPB enzymes (PPAT, PFAS, PAICS, ATIC) identified by SpARC-map to interact with ADSL via ADSL^451-484^, three (PFAS, PAICS, and ATICS) were low/undetectable in non-irradiated controls, but abundant in UV-irradiated samples (Supplementary Data 2). Of the 1717 proteins detected, PFAS, PAICS, and ATIC are among the top 1% of proteins exhibiting statistically significant pulldown enrichment following UV-irradiation, indicating their interaction with ADSL via ADSL^451-484^ is highly specific. PPAT was only detected upon UV-irradiation, but not at statistically significant level. GART, which SpARC-map indicated interacts with ADSL but not via ADSL^451-484^, was abundantly detected, without statistically significant differences, in both UV-irradiated and non-irradiated samples (Fig. 5C-D). These results support SpARC-map’s identification of ADSL^451-484^ as a multispecific interface with multiple, but not all, purinosome enzymes.

### Constraining AlphaFold3 predictions with SpARC-map data

For protein complexes that lack experimentally determined structures, high resolution models from ML-based structure prediction may be able to fill in the gap. Using the PPAT-GART complex as an example, we examined whether PPI interfaces identified by SpARC-map can also be found in AlphaFold3 (AF3) predictions. This interaction was long argued to be biochemically necessary, but direct physical evidence had been lacking. We generated 305 randomly seeded AF3 models of the complex comprised of 4 copies of PPAT, which can exist in both homo-dimeric and homo-tetrameric forms, and 2 copies of GART, which is homo-dimeric. For each model, we identified all atomic contacts (separation <0.4 nm) between PPAT and GART monomers. A majority of AF3 models identified PPAT contacting GART on GART’s AIR synthase (AAs 434-809) and/or GAR transformylase (AAs 809-1010) domains, in various configurations that are not in consensus with each other (Fig. 6A), are structurally incompatible with substrate channeling between PPAT and GART, and are inconsistent with SpARC-map results. Of the regions identified by SpARC-map as PPI interfaces, PPAT AAs 468-504, contacting GART, was found in ~10% of models, while GART AAs 14-55 and 159-232, contacting PPAT, was found in ~2% of the models; only a handful of models possessed PPI interfaces consistent with SpARC-map results (Fig. 6B). Here, SpARC-map results present strong, experimentally grounded constraints on AF3 predictions for the structure of the PPAT-GART complex.

**Figure 6.**
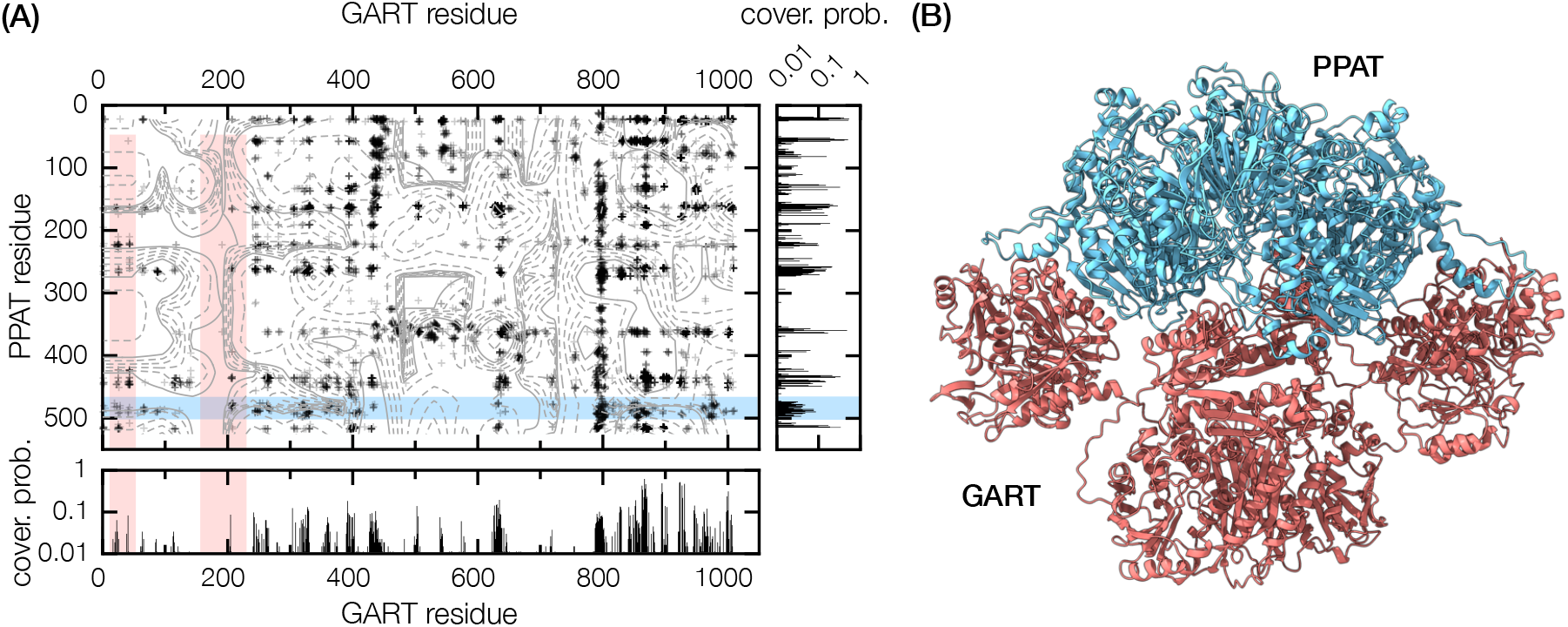
Modeling of the PPAT-GART complex with AlphaFold3. **(A)** Map of atomic contacts between PPAT and GART monomers across 289 AF3 models. Each data point is a PPAT-GART atomic contact found in at least one model; density contours are quadratic interpolations. Also shown are heatmaps indicating the fraction of models that has a specific residue on PPAT (right sub-panel) or GART (bottom sub-panel) to be a part of the PPAT-GART interface. Shaded regions (light blue on PPAT, pink on GART) are regions identified by SpARC-map as probable PPAT-GART interfaces. **(B)** Example of an AF3 model of PPAT-GART complex that is compatible with SpARC-map results; PPAT (homo-tetramer) is show in light blue, GART (homodimer) is in pink.

## Discussion

We have described SpARC-map, a method for identifying the probable PPI interface between two interacting proteins. For p21-PCNA, p53-MDM2, and MYC-MAX, SpARC-map recovered the known PPI interfaces of these complexes with excellent fidelity. For the purinosome, SpARC-map identified hereto unknown PPI interfaces that satisfy structural requirements for substrate channeling, as well multispecific interfaces that participate in multiple PPIs. Using site-specific photocrosslinking in vivo, we validated the identification of ADSL^451-484^ as a multispecific interface via which ADSL interacts with several, but not all, purinosome enzymes. We are now in the process of validating other probable PPI interfaces in the purinosome. Despite being incompletely validated, our SpARC-map data can already provide tantalizing hints into the still obscure cellular mechanisms that regulate purinosome assembly. Several probable purinosome PPI interfaces are decorated with post-transcriptional modifications, e.g. phosphorylations on PPAT T356, GART S625, ADSL Y466, and ATIC T520 (55); these may modulate the strength of PPIs mediated by the decorated interfaces. If true, identifying the responsible modifying enzymes (e.g. kinases) will establish direct connections between cellular signaling pathways and purinosome assembly.

SpARC-map results also provided us new insights into the structural nature of the purinosome. We have previously found C-terminal tagged ADSL is defective at rescuing ADSL-knockout HeLa (Supplementary Materials). This was puzzling because ADSL’s C-terminal region does not contribute to known ligand-binding or active sites (56). Recognizing that this region mediates PPIs between ADSL and multiple purinosome enzymes suggests C-terminal tagging of ADSL disrupts these PPIs, leading to the loss of substrate channeling and possibly destabilizing the purinosome entirely. Given the density of probable direct interactions in the purinosome revealed by SpARC-map, it is unlikely all six enzymes can arrange into a single definite structure that satisfies all binary contacts. This, and the presence of multispecific binding regions, suggest the purinosome is probably not a classic protein complex with an exact stoichiometry. Intrinsically disordered proteins with polyvalent PPIs are known organizers of “fuzzy” complexes such as liquid-like protein condensates (57–60). Polyvalency in structured proteins with multispecific binding sites, e.g. ADSL, further amplified through homo-oligomerization (ADSL is a homo-tetramer), may play a similar role in organizing the purinosome via so-called bridging attractions (61–63).

Presently, SpARC-map can only map PPIs involving bait proteins that express and properly fold in *E. coli*, though split-neo^R^ selection will also function in eukaryote hosts sensitive to the aminoglycoside antibiotic G418. The fragmentation of prey proteins can result in the expression of toxic, unstable, or aggregation-prone peptides (e.g., hydrophobic beta strands) that could preclude split-neo^R^ reconstitution, resulting in biases and a general failure to detect of PPIs mediated by hydrophobic interactions; our SpARC-map implementation, using low-copy plasmids and tunable bait/prey peptide expression, may ameliorate this effect, as could the addition of solubility tags (e.g. SUMO-tag) to further stabilize the prey peptide-neo^R^C fusion. PPI interfaces that require post-translational modifications absent in the *E. coli* host, or require conformation changes (e.g. due to allostery), or are made up of physically proximal but sequentially distal residues, can escape detection; this may be responsible for our failure to detect both reciprocal interfaces for some PPIs. While SpARC-map is able to identify PPI interfaces, it cannot identify contacting residue pairs or definitely constrain the binary complex structure. Lastly, SpARC-map generally fails to detect homo-oligomeric interfaces, because doing so requires prey peptides to outcompete intact proteins for binding, which suppresses their signal. On the other hand, SpARC-map may be especially well-suited for mapping PPIs involving intrinsically disordered proteins/protein domains (IDPs/IDPDs), which are notably common in transcription factors and in the intracellular domains of transmembrane receptors. We expect IDPDs will primarily interact with other proteins via linear peptide sequences, which is the exact scenario SpARC-map was designed to probe.

Aside from direct structural determination, there currently exist mass spectrometry based methods, including hydrogen-deuterium exchange (64), protein painting (65), and crosslinking/mass spectrometry (XL/MS) (66), that can identify PPI interfaces. Compared to those methods, SpARC-map, as a genomics-based method that does not require reconstituting protein complexes in vitro, is technically simpler and instrumentally less demanding, and thus far more accessible. SpARC-map belongs to the class of protein complementation assays (PCAs) (67), including two-hybrid, split-fluorescent protein, and split-enzyme assays which have a long been used for detecting PPIs. Compared to existing PCAs, SpARC-map has two distinct advantages: first, SpARC-map can supply structural information identify both the PPI and the PPI interface; second, the nonspecific background SpARC-map can be directly measured and modelled, allowing its users to make quantitatively rigorous distinction between specific and nonspecific interactions.

SpARC-map will not detect every PPI and PPI interface; for PPIs/PPI interfaces it can detect, SpARC-map results are no substitute for atomic details found in directly determined structures. Nevertheless, given the current BioGRID database curates approximately one million physical interactions in the human proteome (68), if SpARC-map can capture the interface from just a fraction of these, it will constitute a massive expansion in our knowledge of protein complexes. For non-model organisms, where antibody-based assays are often unfeasible (due to the lack of suitable antibodies), SpARC-map, having no reliance on dedicated reagents, can be a tool of first resort for probing PPIs and PPI interfaces in their molecular pathways. Our present view of protein complexes is largely informed by tightly bound complexes with definite structures and stoichiometries, because these are the most amenable to study. Protein complexes bound by dense networks of weak, possible polyvalent PPIs, and which may lack definite structures and stoichiometries, may be far more typical, but have escaped appreciation because they are difficult to probe. SpARC-map, with its sensitivity, is a tool that could enable us to better explore this largely unseen world. Proteins inside a cell never act in isolation; it is their concerted action, with each other and with other biomolecules, that gives the cell life. We believe SpARC-map, given its accessibility and throughput, will be a useful tool when investigating any biological process where PPIs are vital, thereby facilitating discoveries across molecular and cellular biology.

## Materials and Methods

### Structure and design of SpARC vectors

The SpARC vectors pSEP4N and pSEP4C harbor low-copy pBR322 origin of replication and ampicillin-resistance gene. Expression of repressors lacI and tet^R^ (as a single transcript containing an internal ribosome binding site) is controlled by the *lacIq* promoter. Expression of neo^R^N-bait fusion (in pSEP4N) or bait-neo^R^N fusion (in pSEP4C) is controlled by IPTG-inducible *trc* promoter-*lac* operator. The CDS fragment derived peptide coding sequence is inserted upstream of the neo^R^C coding sequence, so that only inserts with no stop codons, and are 3n-tuple in length, can result in the expression of a prey peptide-neo^R^C fusion. Expression of the prey peptide-neo^R^C fusion is controlled by the aTc-inducible *PletO-1* promoter.

### Library construction

Bait CDS is cloned into SpARC vector using the multiple cloning site: p21, PCNA, MDM2^18-125^, GART, PFAS, PAICS, ADSL, and ATIC were cloned as bait into pSEP4N; p53, MYC, MAX, and PPAT were cloned as bait into pSEP4C. Bait-containing SpARC vector is cut at NotI and SbfI sites flanking the peptide library insertion site, blunted, and dephosphorylated. dsDNA containing prey CDS (without stop codon) is digested using DNA Fragmentase (NEB M0348) into 100-300 bp fragments; for multiplexed experiments, prey CDSs are combined in an equimolar mix and fragmented. Prey CDS fragments are purified, blunted, and ligated into the SpARC vector.

### Library pre-selection

The ligated SpARC library is transformed into chemically competent *E. coli* (strain NEBStable) homemade using the TSS-HI method (69), and pre-selected by outgrowing on LB agar containing low levels (2-5 µg/ml) of kanamycin (Supplementary Materials, Table S1). Following outgrowth, colonies are collected, pooled, and the SpARC library is extracted.

### Library final selection

For each pre-selected SpARC library, we measure its kill curve on a series of LB agar plates containing IPTG (various concentrations depending on the bait), aTc (100 ng/mL), ampicillin (100 µg/mL), and varying concentrations of kanamycin (5-640 ug/ml) (Supplementary Materials, Table S1). We also measure the kill curve of the bait-only parental vector under identical conditions. We choose as final selection conditions under which the bait-only SpARC vector cannot survive, but where the SpARC library still yields numerous surviving colonies.

### NGS sequencing

Post final selection, kanamycin-resistant colonies are pooled, collected, and the host SpARC vectors are extracted. The prey peptide coding sequence is amplified using barcoded NGS sequencing primers that anneal on invariant flanking sequences. PCR amplification is for 12-cycles, and the purified PCR product is sequenced using the AmpliconEZ service from Azenta Life Sciences based on Illumina’s MiSeq2 platform, yielding >50k 250 bp paired end reads per sample.

### Prey peptide identification and mapping

Paired-end NGS reads are trimmed, quality-filtered, and merged; gapped sequences are discarded. For each merged NGS read, we identify the longest open reading frame (ORF) that is fused in-frame with neo^R^C, accepting ATG, TTG, GTG as start codons, and map it to prey CDS(s). ORFs that are less than 15 bps in length, or are concatemers (either of CDS fragments from different preys, or of noncontiguous fragments of the same prey), or are unmappable, are discarded. All NGS processing and mapping were done using a custom workflow written in Julia.

### Estimating the nonspecific background

Peptide coding sequences that are inverted, and/or frameshifted with respect to the prey CDS to which they are mapped to, are designated as encoding random peptides. We identify all such random species *i* and quantify their abundances {*n*_*i*_}. The results are fitted to a generalized Pareto distribution (GPD) using maximum likelihood estimation using the Extremes.jl package (25).

### In vivo protein photocrosslinking

HEK293T cells cultured in purine-depleted media were co-transfected with a plasmid expressing 2×StrepTag -ADSL^Y466TAG^, and pMAH-POLY, which expresses the unnatural amino acid incorporation system. Transfected cells were supplemented with 4-azido-L-phenylalanine (1.8 mM) for 48 hours. For photocrosslinking, cells are irradiated using a germicidal UV lamp (260 nm peak emission, 3 minutes) and immediately harvested; non-irradiated cells are collected directly. Cleared cell lysate was incubated with anti-StrepTag magnetic beads (IBA Lifesciences 2-5090-010) and extensively washed. Pulled-down proteins are alklyated and trypsin-digested on-bead. Digested peptides are purified and analyzed via liquid chromatography electrospray ionization tandem mass spectrometry on a Thermo Orbitrap Eclipse. Experiment was repeated with two biological replicates per treatment.

### Mass spectrometry analysis

Peptide are identified and quantified (label-free quantification) using Proteome Discoverer 1.4. To identify differentially enriched protein species, we first estimate the inter-replicate distribution for the abundance of protein species *i, P*_rep_(*n*_*i*_|*n*_*i*, avg_), given its inter-replicate mean *n*_*i*, avg_. We assume *P*_rep_(*n*_*i*_|*n*_*i*, avg_) depends only on *n*_*i*, avg_, and is similar for proteins of similar abundance, i.e. if *n*_*i*, avg_ = *n*_*j*, avg_ = *n*_avg_, then *P*_rep_(*n*_*i*_|*n*_*i*, avg_) = *P*_rep_(*n*_*j*_|*n*_*j*, avg_) = *P*_rep_(*n*|*n*_avg_). We estimate *P*_rep_(*n*|*n*_avg_) by binning together proteins with similar abundances and using Gaussian kernel density estimation, as implemented by the KernelDensity.jl package, to approximate the result. To evaluate whether the abundance of protein species *i* under UV-irradiation is significantly enriched, given its abundance in non-irradiated samples is *n*_*i*, avg, UV-_, we use the estimated *P*_rep_(*n*|*n*_avg_ = *n*_*i*, avg, UV-_) to compute its *p*-value; a small *p* indicates it is enriched in UV-irradiated samples.

## Acknowledgments

ADSL knock-out HeLa was a kind gift from Marie Zikanova, Charles University and General University Hospital, Prague, Czech Republic. We acknowledge the Proteomics and Mass Spectroscopy Core Facility, Huck Institutes for the Life Sciences, Penn State, for the use of their instrumentation. We thank Tony Pedley, Phil Hanoian, Michelle Spiering, Vivek Kapur, Steve Baker, and Han Nguyen for helpful discussions and comments. This work was supported by the U.S. National Institutes of Health under award R01GM024129.

## Supplemental materials

### 1. Schematic of the human de novo purine biosynthetic (DNPB) pathway

**Figure S1.**
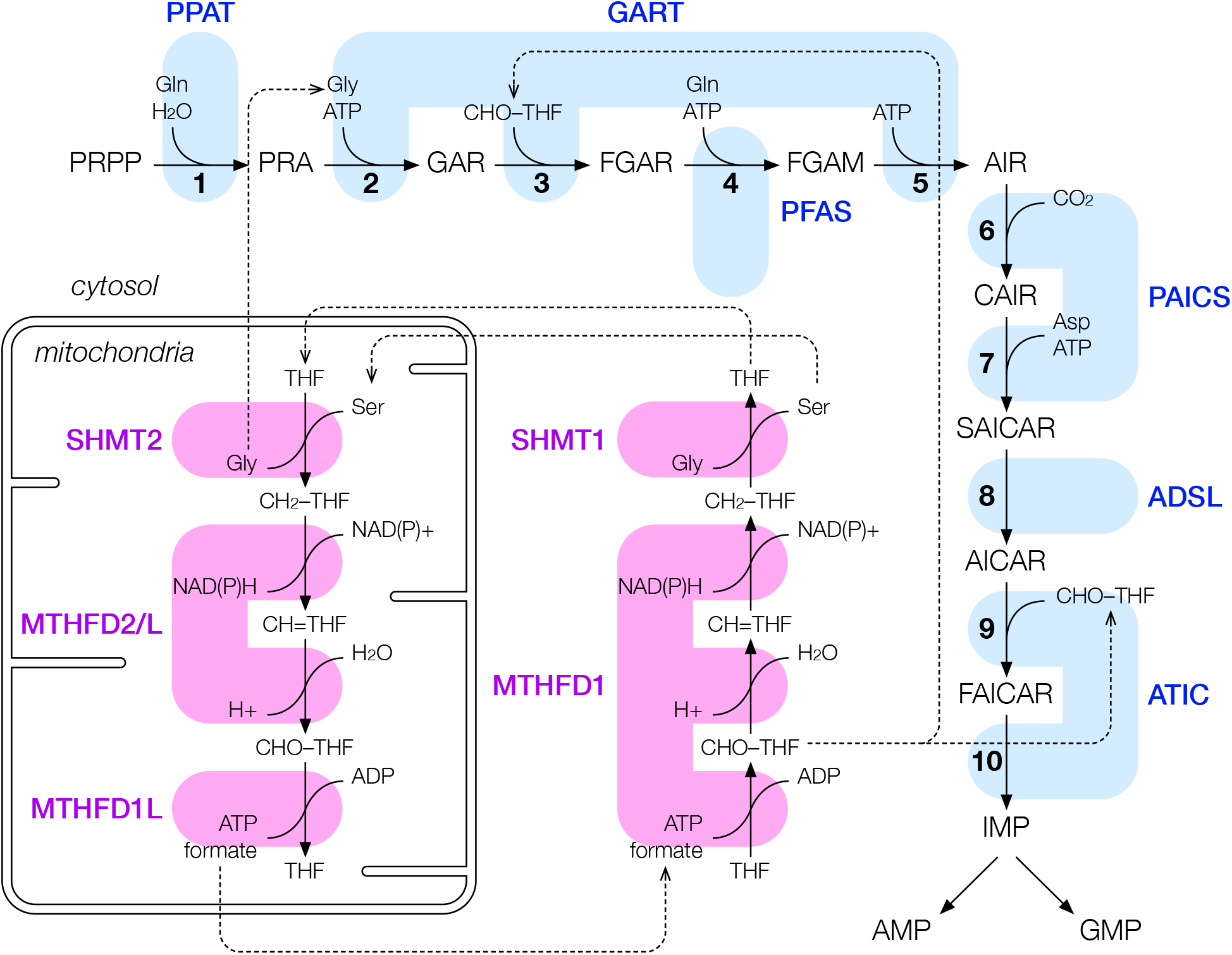
The human DNPB pathway, where 6 DNPB enzymes (blue) convert PRPP (phosphoribosyl pyrophosphate) to IMP (inosine monophosphate) in 10 nonsequential reactions (KEGG: M00048). **Reaction 1: PPAT** (amidophosphoribosyltransferase) converts PRPP to PRA (5-phosphoribosylamine). **Reactions 2, 3: GART** (trifunctional phosphoribosylglycineamide synthetase/formyltransferase/phosphoribosylamino– imidazole synthetase) converts PRA to GAR (glycinamide ribonucleotide) and then to FGAR (phosphoribosyl-N-formylglycineamide). **Reaction 4: PFAS** (phosphoribosylformylglycinamidine synthase) converts FGAR to FGAM (5-phosphoribosylformyl-glycinamidine). **Reaction 5: GART** converts FGAM to AIR (5-aminoimidazole ribotide). **Reactions 6, 7: PAICS** (bifunctional phosphoribosylaminoimidazole carboxylase/succinocarbox– amide synthetase) converts AIR to CAIR (5-phosphoribosyl-4-carboxy-5-aminoimidazole) and then to SAICAR (phosphoribosylaminoimidazolesuccinocarboxamide). **Reaction 8: ADSL** (adenylosuccinate lyase) converts SAICAR to AICAR (5-aminoimidazole-4-carboxamide ribonucleotide). **Reactions 9 and 10: ATIC** (bifunctional 5-aminoimidazole-4-carboxamide ribonucleotide formyltransferase/IMP cyclohydrolase) converts AICAR to

FAICAR (5-formamidoimidazole-4-carboxamide ribotide) and then to IMP. Also shown are one-carbon metabolism enzymes (magenta) responsible for the interconversion between THF (tetrahydrofolate), CH_2_-THF (5,10-methylenetetrahydrofolate), CH=THF (5,10-methenyltetrahydrofolate), and CHO-THF (10-formyltetrahydrofolate) — the last being a necessary cofactor for DNPB. In the mitochondria, these enzymes are **SHMT2** (serine hydroxymethyltransferase), **MTHFD2/2L** (bifunctional methylenetetrahydrofolate de– hydrogenase/cyclohydrolase), and **MTHFD1L** (formyltetrahydrofolate synthetase); in the cytosol, these are **SHMT1** (serine hydroxymethyltransferase), and **MTHFD1** (trifunctional methylenetetrahydrofolate de– hydrogenase/cyclohydrolase/formyltetrahydrofolate synthetase).

## 2. Modeling affinity selection in SpARC-map

Consider a bacterial host in an environment with kanamycin concentration [kan]_out_. The kanamycin concentration [kan]_in_ inside the host cell depends on the balance between kanamycin transport into the cell, and the degradation of kanamycin by reconstituted split-neoR inside the cell,

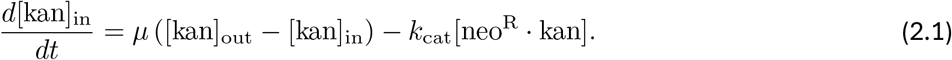

The first term in (2.1) describes kanamycin influx into the cell, characterized by a rate *µ*; the second term describes the degradation of kanamycin by reconstituted split-neoR with catalytic rate *k*_cat_. We assume the inactivation of kanamycin follows Michaelis-Menten kinetics, such that

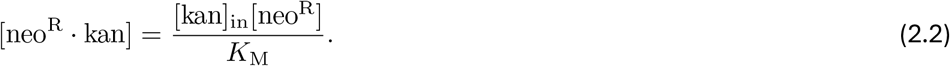

This yields the steady state kanamycin concentration inside the cell, given by

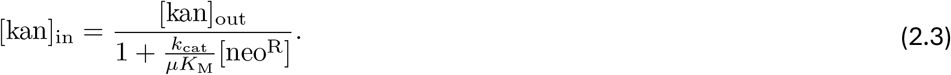

The host can survive only if [kan]_in_ is below some critical value, assumed to be to the minimum inhibitory concentration of kanamycin for *E. coli*, [kan]_MIC_ = 3.4 µM. This imposes a minimum of split-neoR reconstitution,

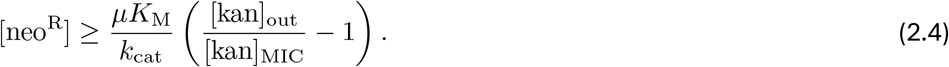

The reconstitution of split-neo^R^ is due to 1:1 bait-prey peptide binding (fused respectively to neo^R^N and neo^R^C). In terms of the apparent dissociation constant *K*_D_^app^, the condition for host survival becomes

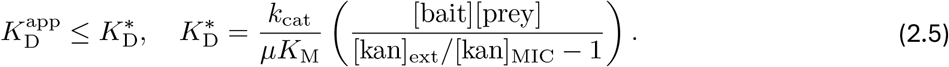

The selection threshold *K*_D_* depends on several kinetic parameters: for the split neo^R^, *k*_cat_/*K*_M_ ≈ 0.1 µM^−1^ s^−1^; for kanamycin transport into *E. coli, µ* ≈ 0.002 s^−1^ (1). Taking [kan]_ext_/[kan]_MIC_ = 5 – 100, and assuming [bait] ~ [prey] = 0.1 – 10 µM (equivalent to ~ 0.01 – 1 mg of a 30 kDa recombinant protein expressed per liter of culture), yields a tunable range for *K*_D_* between 5 nM and 1 mM. This represents the theoretical maximum dynamics range for SpARC-map affinity selection.

**Figure S2.**
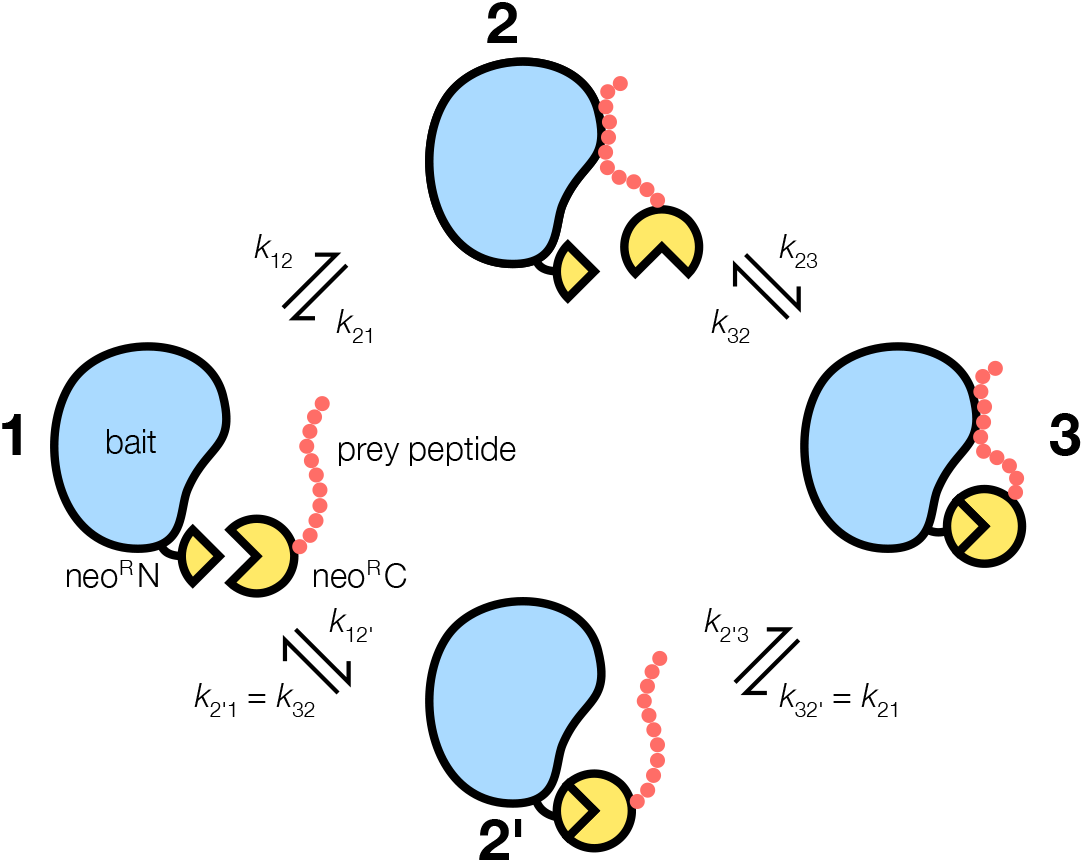
Four-state kinetic model for the reconstitution of split-neoR fragments fused to interacting bait-prey peptide pair. Binding can be initiated either by the bait-prey peptide interaction (state 2), or by the intrinsic interaction between the split-neoR fragments (state 2’). Only states 2’ and 3 contribute to [neo^R^].

There are two contributions to split-neo^R^ reconsitution: the bait-prey peptide interaction (as a transition state), and the intrinsic interaction between neo^R^N and neo^R^C fragments. While bare neo^R^N and neo^R^C fragments do not appear to interact strongly, some intrinsic interaction is probably necessary for reconstituting neoR activity. Assuming the bait-prey peptide and bare neo^R^N-neo^R^C interactions are non-competitive, and a simple 4-state kinetic model (Fig. S2), in the limit bait-prey peptide *K*_D_^PB^ ≪*K*_D_^bare^ between bare neoR fragments, the predominant kinetic path is 1 ↔ 2 ↔ 3, and the reconstitution of split-neo^R^ is given by

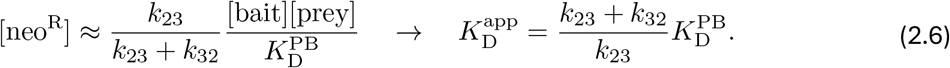

In the opposite limit *K*_D_^PB^ ≫*K*_D_^bare^, the predominant kinetic path is 1 ↔ 2’ ↔ 3, and

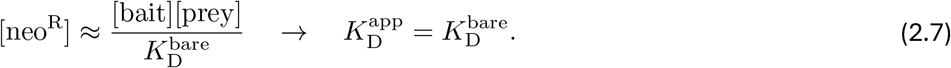

Thus, the real sensitivity limits of SpARC-map affinity selection is limited by the intrinsic interaction between split neo^R^ fragments: when *K*_D_^PB^ ≪*K*_D_^bare^, the apparent dissociation constant *K*_D_^app^ contains a factor (*k*_23_+*k*_32_)/*k*_23_ > 1; when *K*_D_^PB^ ≫*K*_D_^bare^, then affinity between bare neo^R^ fragments represents a sensitivity ceiling.

During selection, even for cells that host the identical bait/prey pair, cell-to-cell variation in bait and prey expression (and therefore split-neo^R^ reconstitution), kanamycin uptake, and kanamycin sensitivity means *K*_D_* is will also vary from cell to cell. The probability of survival *p*_surv_(*K*_D_^app^|*K*_D, avg_*), where *K*_D, avg_* is the average selection threshold across all cells, will be a sigmoidal function with limits *p*_surv_ → 1 when *K*_D_^app^ ≪*K*_D, avg_*, *p*_surv_ → 0 when *K*_D_^app^ ≫*K*_D, avg_* (Fig. S3). For cells transformed with a SpARC library containing many bait-binding prey peptides of varying *K*_D_^app^, characterized by some distribution *P*_lib_(*K*_D_^app^), the NGS species abundance extracted from the post-selection cell population will be directly proportional to *p*_surv_(*K*_D_^app^|*K*_D, avg_*). If selection is weak (large *K*_D, avg_*), it may be difficult to distinguish strong versus moderate binding peptides because their *p*_surv_ will be nearly equivalent. Conversely, strong selection (small *K*_D, avg_*) can drive the library to extinction. Optimal selection is when the decline of *p*_surv_(*K*_D_^app^|*K*_D, avg_*) coincides with the low-*K*_D_ wing of *P*_lib_(*K*_D_). The fraction of transformed cells that survives a given selection condition (characterized by *K*_D, avg_*) is

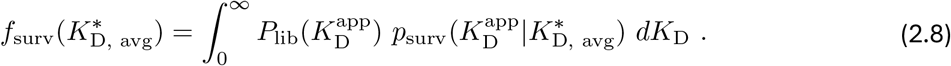

Generically, *f*_surv_(*K*_D, avg_*) will be a sigmoid curve, with limits *f*_surv_ → 1 when *K*_D, avg_* → 0 (no selection) and *f*_surv_ → 0 when *K*_D, avg_* → ∞ (maximum selection).

**Figure S3.**
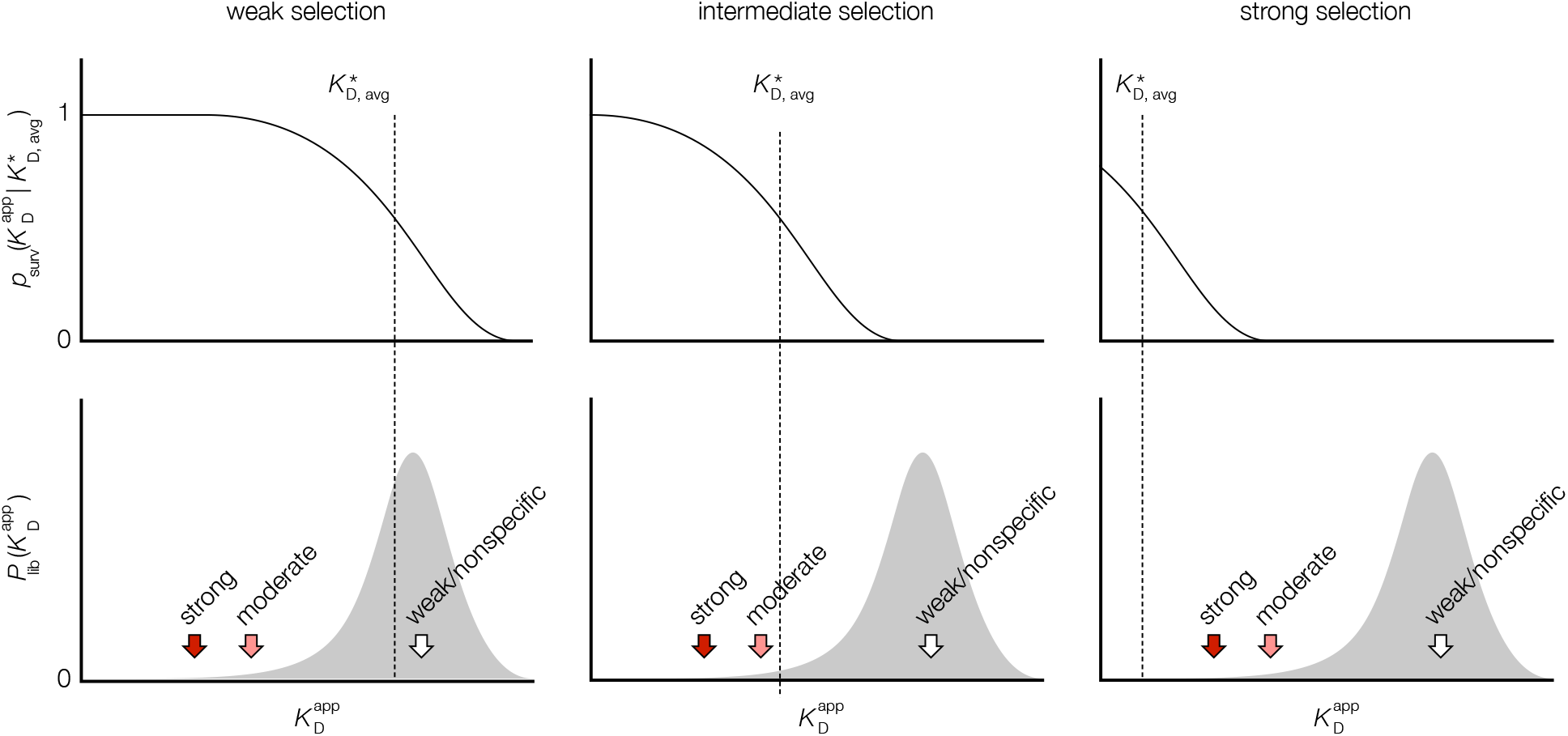
Sketch of affinity selection in SpARC-map. Changing the selection conditions (kanamycin and inducer concentrations) shifts the survival probability *p*_surv_(*K*_D_^app^|*K*_D, avg_*) (top row) to sample different portions of the library affinity distribution *P*_lib_(*K*_D_^app^) (bottom row, showing locations of strong, moderate, and weak/non-specific binders).

## 3. Defective rescue phenotype in C-terminally tagged ADSL

ADSL deficient HeLa (crADSL) requires adenine supplementation in culture media for survival and proliferation (2). We assessed the ability of CMV-driven expression of exogenous ADSL in crADSL to rescue the deficient phenotype by transfecting crADSL with plasmids expressing 2×Strep-ADSL or ADSL-2×Strep fusion. Transfected crADSL was transferred to purine-depleted media and monitored for survival and proliferation. We found crADSL transfected with 2×StrepTag-ADSL had approximately twice as many surviving cells after 2 weeks as crADSL transfected with ADSL-2×StrepTag (Fig. S4A). This suggests ADSL with C-terminal 2×Strep-Tag fusion is defective. After extended culture, both rescue experiments yielded colonies of cells (with many fewer colonies in the ADSL-2×StrepTag) that had re-integrated the exogenous ADSL into the genome; these stably rescued cells can survive and proliferate indefinitely in purine-depleted media. This indicates both 2×StrepTag-ADSL and ADSL-2×StrepTag retained ADSL’s essential enzymatic activity. However, when examined for protein expression, the crADSL::ADSL-2×StrepTag stable rescue expressed (re-integrated) ADSL at ~35× that of wildtype HeLa, while ADSL expression in crADSL:: 2×StrepTag-ADSL is comparable to that of wildtype HeLa (Fig. S4B). Since the C-terminus of ADSL does not participate in known active or ligand binding sites, we interpret the overexpression of ADSL in crADSL::ADSL-2×StrepTag as a compensation for disturbed PPIs involving ADSL-2×StrepTag.

**Figure S4.**
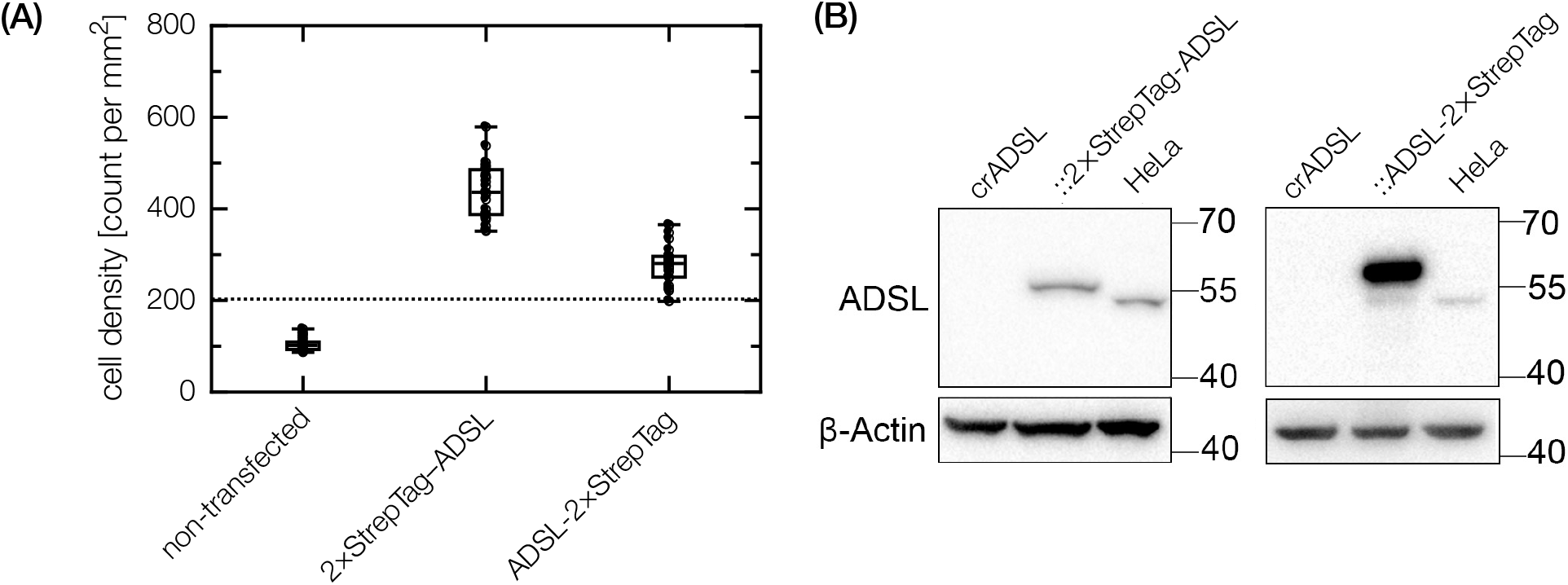
C-terminal tagging of ADSL compromises its ability to rescue ADSL-deficient crADSL cells. **(A)** Proliferation of crADSL rescued by transfection with 2×StrepTag-ADSL or ADSL-2×StrepTag. Dotted line indicates initial crADSL cell seeding density at the start of the rescue assay. Rescue with ADSL-2×StrepTag leads to only half as many surviving cells as rescue with ADSL-2×StrepTag. **(B)** Western blot analysis of reintegrated ADSL expression in crADSL stably rescued with either N-terminally tagged 2×StrepTag-ADSL or C-terminally tagged ADSL-2×StrepTag, showing ADSL hyper-expression is crADSL::ADSL-2×StrepTag.

## Supplementary Tables

**Table S1:** Summary of SpARC-map libraries screened in this study.

| bait | prey | size<br>as ligated<br>($\times 10^3$) | size<br>pre-selected<br>($\times 10^3$) | pre-selection<br>conditions | selection<br>conditions |
| --- | --- | --- | --- | --- | --- |
| p21 | PCNA | 157 | 11 | kan: 4 μg/ml<br>ITPG: none<br>aTc: 100 ng/ml | kan: 40, 56, 80 μg/ml<br>IPTG: none<br>aTc: 100 ng/ml |
| PCNA | p21 | 275 | 26 | kan: 4 μg/ml<br>ITPG: none<br>aTc: 100 ng/ml | kan: 40, 56, 80 μg/ml<br>IPTG: none<br>aTc: 100 ng/ml |
| p53 | MDM2 | 78 | 19 | kan: 4 μg/ml<br>ITPG: none<br>aTc: 100 ng/ml | kan: 7, 14 μg/ml<br>IPTG: 1 mM<br>aTc: 100 ng/ml |
| MDM2<sup>18-125</sup> | p53 | 94 | 20 | kan: 4 μg/ml<br>ITPG: none<br>aTc: 100 ng/ml | kan: 40, 80 μg/ml<br>IPTG: none<br>aTc: 100 ng/ml |
| MYC | MAX (isoform 2) | 60 | 3 | kan: 5 μg/ml<br>ITPG: 1 mM<br>aTc: 100 ng/ml | kan: 14, 20 μg/ml<br>IPTG: 1 mM<br>aTc: 100 ng/ml |
| MAX (iso-<br>form 2) | MYC | 196 | 23 | kan: 4 μg/ml<br>ITPG: none<br>aTc: 100 ng/ml | kan: 20, 40 μg/ml<br>IPTG: none<br>aTc: 100 ng/ml |
| PPAT | PPAT, GART, PFAS,<br>PAICS, ADSL, ATIC | 344 | 51 | kan: 3 μg/ml<br>ITPG: 1 mM<br>aTc: 100 ng/ml | kan: 28, 40, 56 μg/ml<br>IPTG: 1 mM<br>aTc: 100 ng/ml |
| GART | PPAT, GART, PFAS,<br>PAICS, ADSL, ATIC | 61 | 2 | kan: 5 μg/ml<br>ITPG: 1 mM<br>aTc: 100 ng/ml | kan: 28, 40, 56 μg/ml<br>IPTG: 1 mM<br>aTc: 100 ng/ml |
| PFAS | PPAT, GART, PFAS,<br>PAICS, ADSL, ATIC | 188 | 24 | kan: 2 μg/ml<br>ITPG: 1 mM<br>aTc: 100 ng/ml | kan: 20, 28, 40 μg/ml<br>IPTG: 1 mM,<br>aTc: 100 ng/ml |
| PAICS | PPAT, GART, PFAS,<br>PAICS, ADSL, ATIC | 120 | 10 | kan: 5 μg/ml<br>ITPG: : 1 mM<br>aTc: 100 ng/ml | kan: 40, 80, 160 μg/ml<br>IPTG: 1 mM<br>aTc: 100 ng/ml |
| ADSL | PPAT, GART, PFAS,<br>PAICS, ADSL, ATIC | 428 | 35 | kan: 5 μg/ml<br>ITPG: : 1 mM<br>aTc: 100 ng/ml | kan: 40, 80, 160 μg/ml<br>IPTG: 1 mM<br>aTc: 100 ng/ml |
| ATIC | PPAT, GART, PFAS,<br>PAICS, ADSL, ATIC | 92 | 8 | kan: 5 μg/ml<br>ITPG: : 1 mM<br>aTc: 100 ng/ml | kan: 40, 80, 160 μg/ml<br>IPTG: 1 mM<br>aTc: 100 ng/ml |

Note: cell outgrowth for preselection and final selection at 30°C.

**Table S2.**
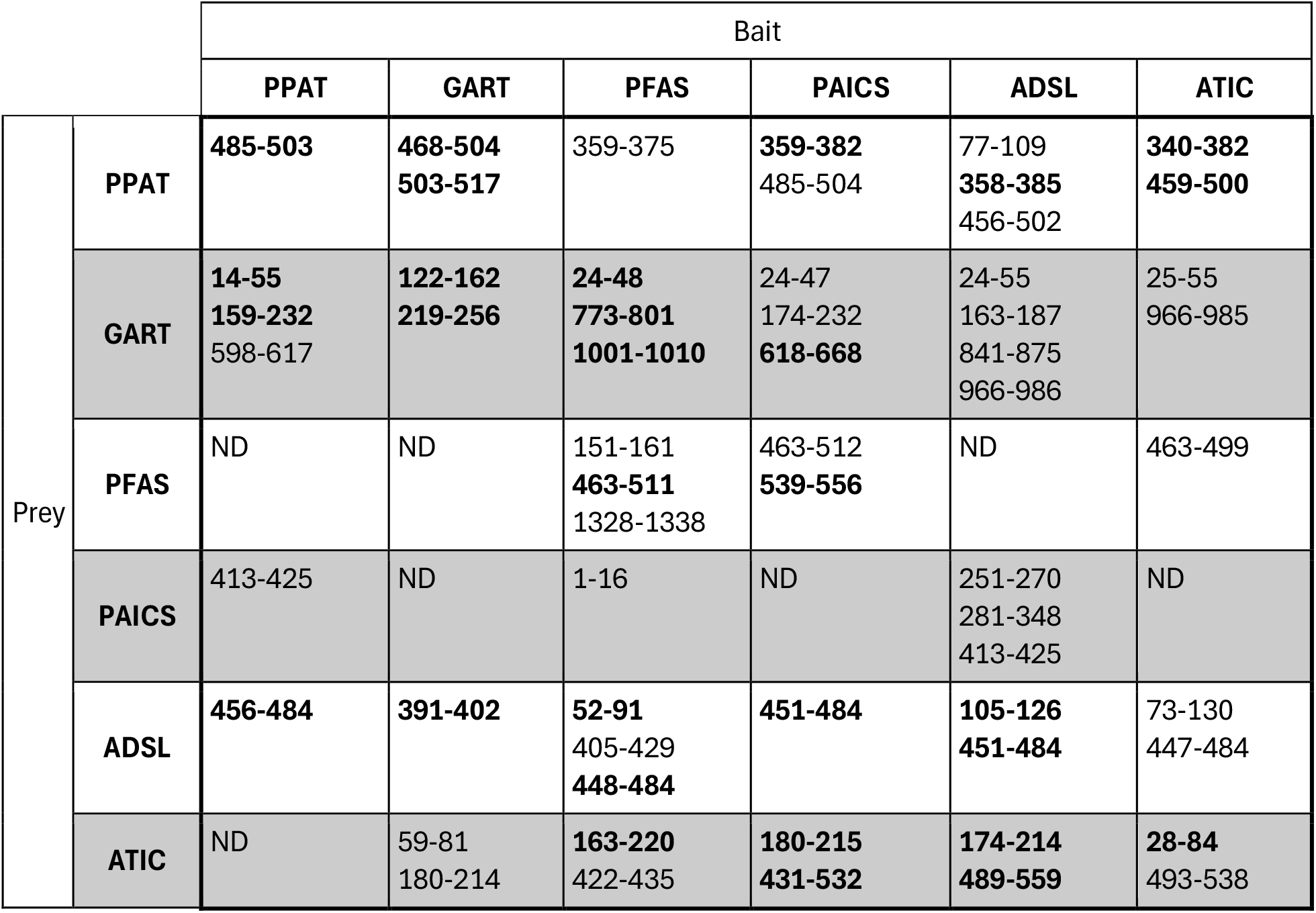
Probable PPI interfaces in the purinosome identified by SpARC-map (*p*<0.01, bold *p*<0.003)

## Notes

### Competing Interest Statement

A provisional patent has been filed with Jingxuan He, Ling-Nan Zou, and Stephen J. Benkovic as inventors.

### Summary of Updates

Includes two additional examples (p53-MDM2, MAX-MYC) of recovering known PPI interfaces using SpARC-map.

